# Organelle-targeted Laurdans measure heterogeneity in subcellular membranes and their responses to saturated lipid stress

**DOI:** 10.1101/2024.04.16.589828

**Authors:** Adrian M. Wong, Itay Budin

## Abstract

Cell organelles feature characteristic lipid compositions that lead to differences in membrane properties. In living cells, membrane ordering and fluidity are commonly measured using the solvatochromic dye Laurdan, whose fluorescence is sensitive to membrane packing. As a general lipophilic dye, Laurdan stains all hydrophobic environments in cells, so it is challenging to characterize membrane properties in specific organelles or assess their responses to pharmacological treatments in intact cells. Here, we describe the synthesis and application of Laurdan-derived probes that read out membrane packing of individual cellular organelles. The set of Organelle-targeted Laurdans (OTL) localizes to the ER, mitochondria, lysosomes and Golgi compartments with high specificity, while retaining the spectral resolution needed to detect biological changes in membrane packing. We show that ratiometric imaging with OTL can resolve membrane heterogeneity within organelles, as well as changes in membrane packing resulting from inhibition of lipid trafficking or bioenergetic processes. We apply these probes to characterize organelle-specific responses to saturated lipid stress. While ER and lysosomal membrane fluidity is sensitive to exogenous saturated fatty acids, that of mitochondrial membranes is protected. We then use differences in ER membrane fluidity to sort populations of cells based on their fatty acid diet, highlighting the ability of organelle-localized solvatochromic probes to distinguish between cells based on their metabolic state. These results expand the repertoire of targeted membrane probes and demonstrate their application to interrogating lipid dysregulation.

## Introduction

The lipid membranes that compartmentalize eukaryotic cells and their organelles are diverse biological structures that vary dramatically in composition, topology, and function. Membrane fluidity, which describes the lateral mobility of lipids and membrane proteins within the lipid bilayer (*1*), and other associated properties (*2*), modulate processes in metabolism, signaling, trafficking, migration, and exo/endocytosis (*3–5*). Several facets of lipid chemistry broadly control membrane fluidity through their effects on bilayer packing, including acyl chain length and saturation, head group chemistry, and cholesterol content (*6–8*). Organisms use these lipidome features to regulate membrane fluidity; for example, in response to environmental changes (*9–11*). However, metabolic disorders can also cause dysregulation of membrane fluidity, commonly by increasing levels of saturated fatty acids and cholesterol (*12–16*). For example, erythrocytes from diabetic patients feature decreased membrane fluidity that can impair circulation (*17*), while liver fibrosis causes reduced hepatocyte membrane fluidity whose reversal can rescue disease pathology (*14*). Excess saturated fatty acids are incorporated by cells into complex lipid species such as ceramides and disaturated glycerolipids, which modulate endogenous cell signaling (*18*, *19*) and induce stress response pathways (*20*, *21*). While the incorporation of saturated fatty acids into cell membranes has long been proposed to act on specific organelles (*22*), it has been challenging to resolve their differential effects on specific intracellular membranes. Therefore, new tools are required to study specific sites of membrane fluidity dysregulation.

A variety of amphiphilic solvatochromic dyes that readily incorporate into lipid bilayers have been employed to study membrane fluidity in synthetic liposomes, as well as intact cells and tissues (*23–25*). Among these, 6-lauroyl-2-dimethylamino naphthalene (Laurdan) has been the most widely used probe due to its high sensitivity to changes in membrane packing and phase transition, strong association with lipid environments in cells, and its emission in the blue/green spectra, which allows for co-labeling with fluorophores that emit in red and far red regions (*26–28*). Upon excitation, Laurdan exists within either a locally excited state native to the fluorophore, or an internal charge transfer state due to a larger dipole moment (*29*). In the internal charge state, Laurdan reorients the dipole of neighboring water molecules, resulting in energy loss to the surrounding environment and ensuing a bathochromic (red) shift in its emission (*30*). In more fluid membranes, the degree of membrane water penetration is greater due to the loosely packed lipids, leading to greater dipolar relaxation and a more dramatic shift in its emission (*26*, *30*). Therefore, Laurdan is sensitive to the fluidity of cell membranes, and can resolve both bulk differences in fluidity between membranes and local micrometer domains within membranes (*26*, *31–33*). Laurdan emission changes can be read out with fluorometry or microscopy and differences in fluidity are often quantified via the normalized difference between emission intensity in two emission wavelengths of spectral windows (channels), corresponding to blue and green emission regions, in a value known as generalized polarization (GP) (*34*). Alternative modalities, including spectral deconvolution using linear array detectors or fluorescence lifetime imaging (FLIM) (*35*, *36*), can provide additional resolution and details on the underlying membrane packing read out by Laurdan.

While Laurdan is a powerful tool for studying membrane biophysics, it rapidly internalizes, leading to staining of all membrane organelles (*27*). The overall density of membrane organelles within a cell and high degree of contact makes visualizing organelle specific changes in membrane fluidity difficult (*30*, *37*, *38*). Typically to overcome this issue, an organelle marker is used as a mask in conjunction with dye in multi-channel confocal microscopy (*39*, *40*). However, the close proximity of many organelles within cells is far below that of the resolution of light microscopy, making interpretation of observed heterogeneities in organelle membrane fluidities challenging to interpret. The popular Laurdan derivative, C-Laurdan, in which a methyl group on the dimethylamine of Laurdan is modified to a carboxylic acid, features improved spectral properties and better stains the plasma membrane (PM) than Laurdan. C-Laurdan still features a high degree of internalization, which can be prevented by additional modification with charged groups, effectively labeling the outer leaflet of the PM (*27*). Recently, intracellular organelle targeted probes have been developed in which small chemical moieties that target accumulation of solvatochromic dyes to specific intracellular chemical environments have been developed. These include 1) Flipper probes that accumulate at the PM or one of three organelles and read out membrane tension but also lipid packing by FLIM (*41*) 2) variants of the solvatochromic dye Nile Red that are targeted to a range of organelle compartments and spectrally respond to mechanical and osmotic stress (*38*) and 3) a molecular rotor (Mitorotor-1) that also accumulates in the inner mitochondrial (IMM) and reports changes in fluidity by FLIM in response to electron transport chain (ETC) activity (*42*). However, no such equivalents have been developed based on Laurdan or related variants, nor have the capacity for such probes to read out organelle-specific lipidome responses been tested.

Here, we report a set of organelle targeted probes based on C-Laurdan that label ER, Golgi, mitochondrial membranes. We show that these organelle targeted Laurdans (OTLs) localize to their respective organelles in HeLa cells and retain sensitivity to membrane packing. We use OTLs to report the distribution of apparent membrane fluidity in organelles and how they respond to inhibition of intracellular lipid transport. We then show that OTLs are effective tools for monitoring organelle-specific changes in response to exogenous saturated lipid stress.

## Results and Discussion

### Synthesis and validation of Organelle-targeted Laurdans

For each OTL, we modified C-Laurdan through conjugation of its carboxylic acid with primary amine-containing chemical moieties via HATU coupling (Figure S1). ER-Laurdan (Figure 1A) contains a pentafluorophenyl (PFP) group with a PEG_10_ linker; the PFP group reacts with thiols in unfolded proteins in the ER and the linker provides flexibility for the protein-bound probe to integrate into the membrane (*43*, *44*). Golgi-Laurdan (Figure 1B) mimics the structure NBD C6 Ceramide, a dye that accumulates in the Golgi via its trafficking from the ER via the ceramide transporter CERT (*45*). To maintain the orientation of Laurdan, with the Prodan fluorophore near the hydrocarbon interface of the bilayer, we coupled Sphingosine (d12:1) base directly with the carboxylic acid of C-Laurdan. Lyso-Laurdan (Figure 1C) accumulates in the lysosome via a morpholine group that is entrapped in lysosome via protonation to its cationic form in the acidic compartment (*41*). Mito-Laurdan (Figure 1D) contains a cationic triphenylphosphonium moiety, which accumulates at the inner mitochondrial membrane due to its negative membrane potential, connected via a 3 carbon linker (*46*). Synthesis and purification of each compound is described in the Materials and Methods section.

**Figure 1.**
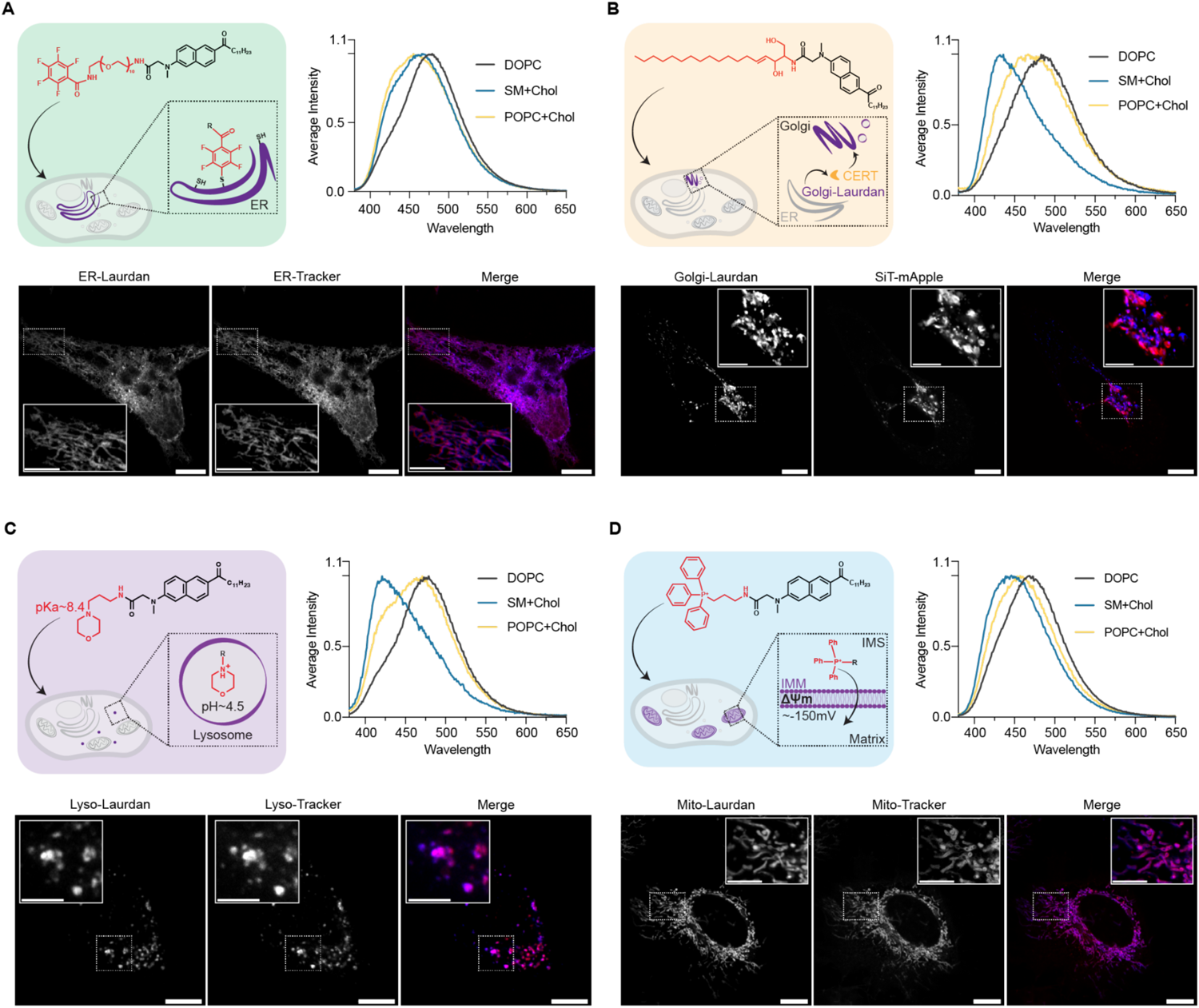
Mechanism of targeting and validation of four OTLs. **A**. ER-Laurdan contains a PFP group that reacts with free thiols in the ER, tethering near lumenal membranes. Solvatochromic properties of the dye were assessed in DOPC, SM+Chol and POPC+Chol vesicles and a hypochromic shift in emission in the SM+Chol and POPC+Chol vesicles. Localization of the dye was assessed by confocal microscopy using ER-Tracker Red and showed high localization of ER-Laurdan to the ER. Insets highlight ER tubules. Scale bars, 10 µm or 5 µm insets. **B**. Golgi-Laurdan mimics the structure of ceramide, which is trafficked to the Golgi. A bathochromic shift in DOPC vesicles compared to SM+Chol and POPC+Chol vesicles as expected. Localization was assessed with the Golgi anchored SiT-mApple. **C.** Lyso-Laurdan contains a morpholino group that is protonated to its cationic form in acidic compartments, trapping in the lysosomal lumen. Spectral shifts occurred as expected between vesicle populations. Localization of the dye was assessed with Lyso-Tracker Deep Red, showing high overlap between the two channels. **D**. Mito-Laurdan contains a TPP group that is drawn to the mitochondrial matrix by the negative IMM potential. Spectral shifts between vesicle populations were as expected. Localization of Mito-Laurdan was assessed by costaining with Mito-Tracker Deep Red indicated high localization of Mito-Laurdan to mitochondria.

We first tested whether replacing the carboxylic acid of C-Laurdan with an amide linkage to each of the four targeting groups disrupts Laurdan’s solvatochromic properties. Each OTL retained the absorption spectra of the parent compound, as well as its low fluorescence in aqueous buffers (Figure S2). They also retained solvatochromic behavior in solvents: a progressive blue-shift as polarity decreases (DMSO vs. ethanol vs. chloroform) (Figure S3). We then incorporated each compound into large unilamellar vesicles (LUVs) composed of either 100 mol % 1,2-dioleyol-sn-glycerol-3-phosphatidylcholine (DOPC), 55 mol % 1-palmitoyl-2-oleoyl-glycero-3-phosphatidylcholine (POPC) + 45 mole % cholesterol (Chol), or 55 mol % sphingomyelin (SM) + 45 mol % Chol. Bilayer packing for these compositions spans a range from highly disordered (DOPC) to highly ordered (SM/Chol). Each OTL showed increased blue-shifted emission (400-420 nm) and reduced red-shifted emission (490-510 nm) in SM/Chol and POPC/Chol vesicles compared to DOPC vesicles, consistent with a retention of solvatochromic responsiveness (Figure 1A-D). Quantified as Generalized Polarization (GP) values, these shifts were similar to that of Laurdan and C-Laurdan for that of Lyso- and Golgi-Laurdans, but reduced in magnitude for ER- and Mito-Laurdans at both 25 °C (Figure S4) and 37 °C (Figure S5). This diminished sensitivity could result from the attachment of the polar groups – a PEG_10_ linker in ER-Laurdan and phosphonium in Mito-Laurdan – pulling electron density from the methylamine of Laurdan, reducing the formation of the charge transfer state required for dipolar relaxation. However, even these probes showed clear emission shifts in both liposomes and whole cells, as described below.

We next assessed the localization of the OTLs in HeLa by confocal microscopy, comparing the pattern of their blue emission with that of red far-red organelle markers. ER, mitochondrial, and lysosomal Laurdans showed high signal overlap between commonly used organelle stains ER Tracker, MitoTracker Far Red, and LysoTracker Far Red, respectively (Figure 1A,C, D). Imaging with ER-Laurdan was able to stain ER tubules that are typically difficult to resolve from other organelles in close proximity by general Laurdan staining. For the Golgi, we also observed signal overlap for Golgi-Laurdan to the *cis*-Golgi marker sialyltransferase (SiT)-mApple, both appearing as compact perinuclear structure with moderate ER staining. There were some differences in the localized distributions of the dyes within the Golgi apparatus, potentially reflecting differences in cisternal distributions of SiT and ceramides within the Golgi. Compared to NBD C6 Ceramide, Golgi-Laurdan accumulation also required an extended 24-hour incubation post-staining. We suspect the slow trafficking of Golgi-Laurdan is due to its relative bulkiness and the proximity of Laurdan’s naphthalene ring to the ceramide headgroup, which could contribute to weakened interaction with CERT (*47*, *48*).

### Characterization of intra-organelle membrane heterogeneity

We next asked if OTLs could measure heterogeneity in membrane fluidity within individual organelles, which are challenging to resolve by non-targeted solvatochromic probes due to the close proximity of intracellular compartments. After staining HeLa cells with each OTL, we acquired emission in ordered and disordered spectral windows simultaneously by confocal microscopy to calculate GP at each pixel; the resulting distribution is shown with heat maps and frequency histograms in Figure 2A-D. We hypothesized that GP distributions within organelles would reflect both the underlying membrane heterogeneity and the intrinsic sensitivity of each probe to membrane packing, as characterized in model membranes.

**Figure 2.**
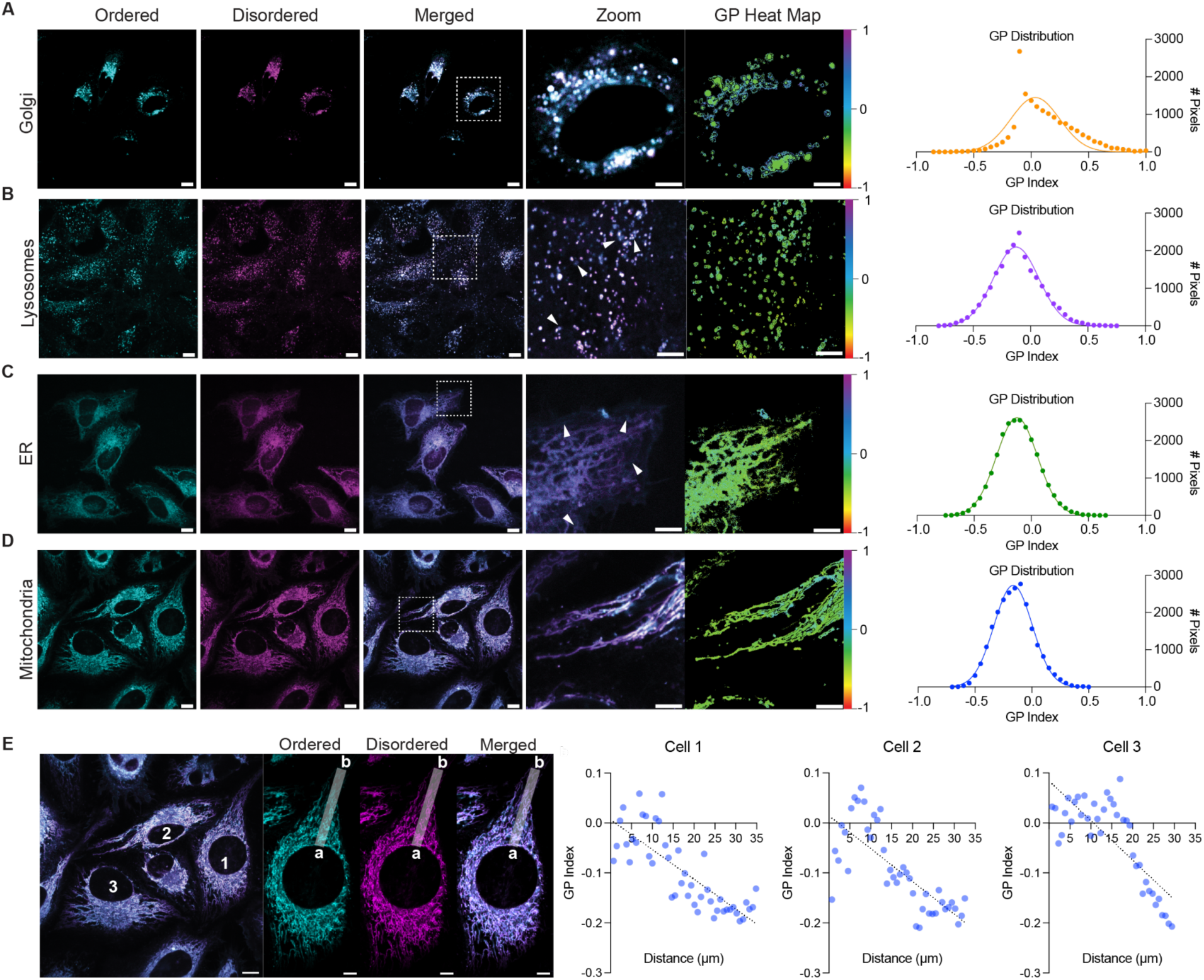
OTLs identify robust subcellular heterogeneity. **A**. Cells stained with Golgi-Laurdan, ordered and disordered channels are shown on the left two panels with merged images to the right. Insets and corresponding GP heatmap highlight heterogeneity within the endomembrane system, with an ordered perinuclear Golgi. Scale bars, 10 µm or 5 µm for inset. Histogram distribution of the GP from each pixel from the inset is fitted to a Gaussian Function with an R^2^ = 0.7278. SD from the mean GP is 0.250. **B.** Cells stained with Lyso-Laurdan reveal differences in fluidity between lysosomes. Insets and corresponding GP heatmap highlight a range of ordered (arrows) and disordered compartments. Scale bars, 10 µm or 5 µm for inset. Gaussian fit of the histogram has an R^2^ = 0.9816. SD from the mean GP is 0.208. **C**. Cells stained with ER-Laurdan show membrane ordering across ER tubules. Insets and corresponding GP heatmap highlight ordered regions that are at the peripheral of the network near the PM (arrows). Scale bars, 10 µm or 5 µm for insets. Histogram distribution fitted to a Gaussian function has an R^2^ = 0.9985. SD from the mean GP is 0.180, indicating a lower degree of heterogeneity than for Golgi and lysosomes. **D**. Cells stained with Mito-Laurdan show the lowest overall heterogeneity, with a Gaussian fit of R^2^ = 0.9932 and SD of GP of 0.167. Insets highlight peripheral mitochondria with higher disordered channel intensities. Scale bars, 10 µm or 5 µm for insets. **E**. Quantified changes in GP between perinuclear mitochondria and peripheral mitochondria are shown for three cells. An example of the plot profile used to calculate the GP trend is shown to the right from point *a* (nuclear mitochondria) to point *b* (peripheral mitochondria). The calculated GPs were fitted to a simple linear regression, showing a correlation between distance from the nucleus and decreasing GP/membrane ordering.

Among the four organelles tested, only the Golgi and lysosomes feature substantial abundances of cholesterol and sphingolipids, well-established modulates of Laurdan GP. As an intermediate secretory compartment, Golgi membranes have been proposed to be characterized by lipid heterogeneity that could be relevant for membrane trafficking (*49*). Among OTLs, Golgi-Laurdan also exhibited the highest GP heterogeneity, with a standard deviation (SD) of GP equal to 0.250. We observed that perinuclear Golgi regions featured a lower GP (Figure 2A). Low GP regions could thus reflect an enrichment of *cis*-Golgi cisternae, which feature reduced abundances of cholesterol and sphingomyelin (*50*, *51*). Lyso-Laurdan also showed high heterogeneity (SD = 0.208), with distinct compartments showing elevated GP states (Figure 2B). When cells are grown on serum, cholesterol is taken up by endosomes through the endocytic pathway and then cholesterol efflux from the lysosomes to the rest of the cell. Because the pH-sensitive morphiline group localizes Lyso-Laurdan to both late endosomes and lysosomes, this heterogeneity could thus represent multiple endo-lysosomal compartments or lysosomes in different states of cholesterol trafficking (*52*).

In contrast to lysosomes and the Golgi, ER and mitochondrial fluidity appeared more uniform across cells, which is consistent with the predominance of unsaturated phospholipids in these compartments. However, we could still observe regions of higher ordering in ER tubules, often extending to the PM, despite the overall lower GP heterogeneity of ER-Laurdan (SD = 0.180), (Figure 2C). Mito-Laurdan featured the lowest GP heterogeneity of the OTLs tested in HeLa cells (SD = 0.167), but we could still characterize regions with lower GP, which corresponded to peripheral mitochondria (Figure 2D). Within single cells, we measured a consistent decrease in Mito-Laurdan GP with distance from the nucleus (Figure 2E). Given the dependence of IMM packing on membrane potential, described further below, this trend could reflect increased membrane potential of peripheral vs. perinuclear mitochondria that has been previously observed across several cell types (*53*).

### Pharmacological modulation of lysosome and mitochondrial membranes

We hypothesized that membrane fluidity read out by OTL would be sensitive to organelle-specific changes in lipid composition. The lysosome membrane-localized Niemann-Pick type C1 protein (NPC1) serves as a key mechanism for cholesterol efflux to other organelles, so inhibition of NPC1 leads to cholesterol accumulation in lysosomes (*52*). To assess whether increased cholesterol in the lysosomes would lead to decreased membrane fluidity, we treated HeLa cells with the NPC1 inhibitor U1866A (*54*) and read out changes in GP using Lyso-Laurdan. We observed that NPC1 inhibition caused the heterogeneous lysosomal population to become more uniformly ordered (SD = 0.191), with an overall increase in GP that was consistent with higher membrane cholesterol content (Figure 3A). Thus, differences in cholesterol efflux could drive heterogeneity in lysosomal membranes.

**Figure 3.**
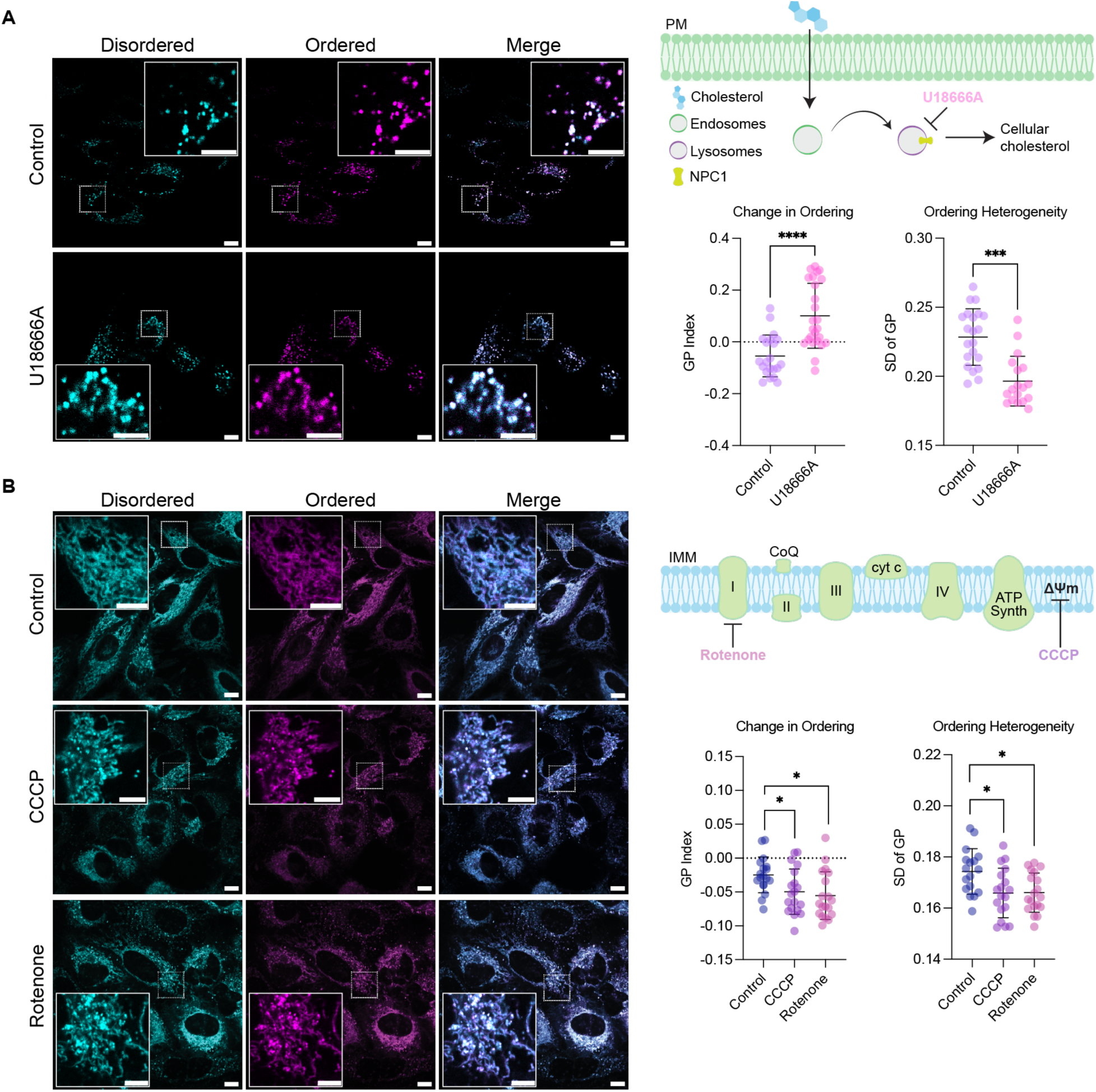
Chemical modification of organelle membrane fluidity. **A.** HeLa cells stained with Lyso-Laurdan were treated with 10 µM U18666A, which inhibits the distribution of cholesterol from lysosomes to other organelles by inhibiting NPC1. U18666A-treated cells show an increase in GP, indicating more ordered membranes, and a reduction in GP heterogeneity as measured by its SD within individual cells. Representative images are shown in the left panel with insets highlighting ordered, swollen lysosomes in U1866A-treated cells compared to smaller, more disordered, and heterogenous lysosomes in control cells. Scale bars, 10 µm or 5 µm (inset). **B.** HeLa cells were stained with Mito-Laurdan and then treated with the uncoupler CCCP, which ablates membrane potential 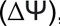 , or Rotenone, which inhibits Complex I. Both compounds lead to a decrease in GP, indicating a more fluid IMM, and GP heterogeneity. Representative images are shown in the left panel with insets highlighting transition from tubular, ordered mitochondria in control cells to fragmented, disordered mitochondria in uncoupled cells. Error bars represent ±SD; *n* = 20 cells per condition. *p<0.05; ****p<0.0001.

We also tested whether Mito-Laurdan could detect changes in IMM fluidity in response to changes in respiratory state. It was recently reported that uncoupling of the ETC led to increased inner membrane fluidity using the mitochondrially-localized molecular rotor probe, Mitorotor-1 (*42*). To determine whether Mito-Laurdan could similarly detect changes in fluidity caused by uncoupling of respiration, we treated HeLa cells with the uncoupler carbonyl cyanide m-chlorophenyl hydrazone (CCCP). Mito-Laurdan remained localized to mitochondria upon CCCP treatment but showed a decrease in GP, suggesting an increase in membrane fluidity upon uncoupling. It was previously proposed that uncouplers could increase fluidity by promoting electron transfer reactions (*42*). To test this, we also applied the Complex I inhibitor, Rotenone, which directly reduces electron flux through the ETC. Rotenone treatment acted similarly to CCCP, reducing membrane ordering, which was also previously observed with the Complex III inhibitor Antimycin A (*42*). These experiments suggest that changes in apparent IMM ordering could result from changes to membrane potential itself, though we also cannot rule out a localized redistribution of potential-sensitive dyes in response to membrane potential. For both CCCP and Rotenone treatment, ordering heterogeneity (SD of GP) also decreased, suggesting that metabolic differences between mitochondria could contribute to the intracellular membrane heterogeneity we and others have observed.

### OTLs distinguish organelle-specific responses to saturated fatty acids

We next asked if OTLs could detect organelle-specific changes in fluidity due to metabolic dysregulation. Palmitic acid (PA) is the most common mammalian saturated fatty acid (*55*) and elevated intracellular PA concentrations have been implicated in numerous diseases(*22*, *56*, *57*). Treatment of cells with PA is thus a common model for saturated lipid stress associated with metabolic disorders. We incubated HeLa cells with Bovine Serum Albumin (BSA)-complexed PA in concentrations up to 200 μM, a range in which all organelles retained regular morphology (Figure S6). ER-, Golgi-, and Lyso-Laurdans all showed an increase in GP for 200 μM PA, indicating reduced membrane fluidity (Figure 4). Lyso-Laurdan also showed a GP increase at the moderate 100 μM treatment, suggesting increased saturated lipid incorporation into lysosomal membranes and/or increased sensitivity of the Lyso-Laurdan probe. The Golgi showed complex responses to PA, with an initial decrease in GP for 50 µM treatment. Only Mito-Laurdan GP was unaffected by 200 μM PA, suggesting that the IMM is more resistant to changes in lipid packing induced by saturated lipid stress. This finding is consistent with our previous data in yeast, where mitochondrial membrane composition was less sensitive to increases in saturated lipids compared to the rest of the cell (*58*).

**Figure 4.**
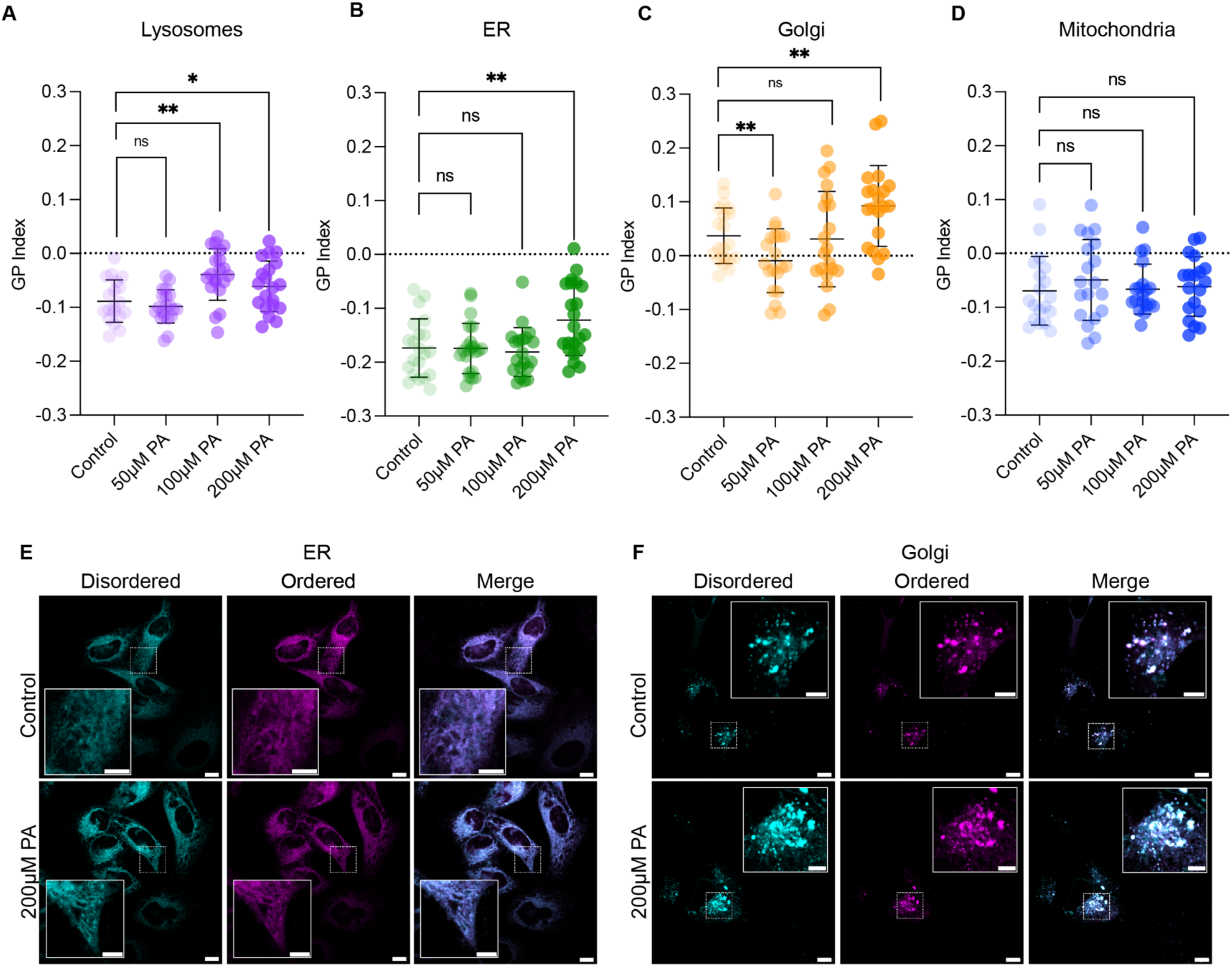
OTLs differentiate organelle specific changes to saturated fatty acid stress. HeLa cells were treated with 50 µM, 100 µM or 200 µM PA or a BSA vehicle control then stained with OTLs. **A**. Cells stained with Lyso-Laurdan that were treated with 100 µM and 200 µM PA showed a significant increase in GP, indicating more ordered lysosomes compared to control. Error bars represent ±SD. ns P>0.05; * p<0.05; ** p<0.01. **B**. ER-Laurdan stained cells showed no significant change in GP at 50 µM and 100 µM PA treatment but a significant increase in GP at 200 µM PA from control. Cells stained with Golgi-Laurdan showed a decrease in GP at 50 µM PA treatment, indicating a more fluid Golgi with low concentrations of exogenous saturated fatty acids. There was no change in GP between control and 100 µM PA treated cells but a significant increase with 200 µM treatment compared to control. **D**. Mito-Laurdan stained cells showed no difference in GP between treatment groups. **E**. Representative images from control and 200 µM PA treated cells stained with ER-Laurdan. Insets highlight more ordered ER tubules in the PA treated group compared to the more fluid control tubules. Scale bars, 10 or 5 µm (inset). **F**. Representative images from Golgi-Laurdan stained control and 200 µM PA treated cells. Insets highlight more ordered Golgi with 200 µM PA treatment. Images for the rest of the organelles and PA concentrations are shown in Figure S6.

### Sorting of cells based on their lipid diets using ER-Laurdan

Based on the ability of specific OTLs to read out changes in membrane fluidity resulting from PA treatment, we hypothesized that these probes could allow for evaluation and isolation of cells with differing lipid metabolism based strictly on membrane biophysical properties. We chose to utilize ER-Laurdan, which has modest sensitivity in synthetic vesicles compared to other OTLs, but targets an organelle that acts as a hub for lipid metabolism and saturated lipid stress (*21*). We first tested the sensitivity of ER-Laurdan GP on a population of detached cells, finding that bulk emission was sensitive to PA treatment across a range of temperatures (Figure S7). The emission spectral changes in cells was generally larger than that in liposomes, suggesting that the lack of targeting and retention in the latter could have contributed to its reduced apparent sensitivity we initially observed.

We asked if cells could be sorted based on their lipid diet using fluorescence of ER-Laurdan (Figure 5). We carried out a metabolic labeling experiment using the deuterated PA palmitic acid-d2 (PA-d2). After a 2-day treatment with 200 µM PA-d2 or a BSA vehicle control, cells were detached, mixed, stained with ER-Laurdan, and sorted using a blue filter and green (AmCyan) filter for the ordered and disordered emissions, respectively. The mixed cell population was sorted by FACS into three groups, P5 which had high DAPI:AmCyan ratio (ordered), P6 with a low ratio (disordered), and an intermediate population P4 (Figure S8). Sorted cells were then analyzed for PA-d2 incorporation by fatty acid methyl esters (FAME) analysis, with the incorporation of PA-d2 unambiguously identified with Mclafferty Ions with an *m/z* of 76 vs. *m/z* of 74 for unlabeled fatty acids. As expected, deuterated FAMEs were concentrated in the highly ordered P5 population, reduced in the intermediate ordering population P4, and were largely absent from the disordered population P6, corresponding to differing ratios of PA-d2 fed to unfed cells. This experiment indicates that cell sorting based on spectral shifts in ER-Laurdan is sufficient to isolate cells based on differences in lipid incorporation.

**Figure 5.**
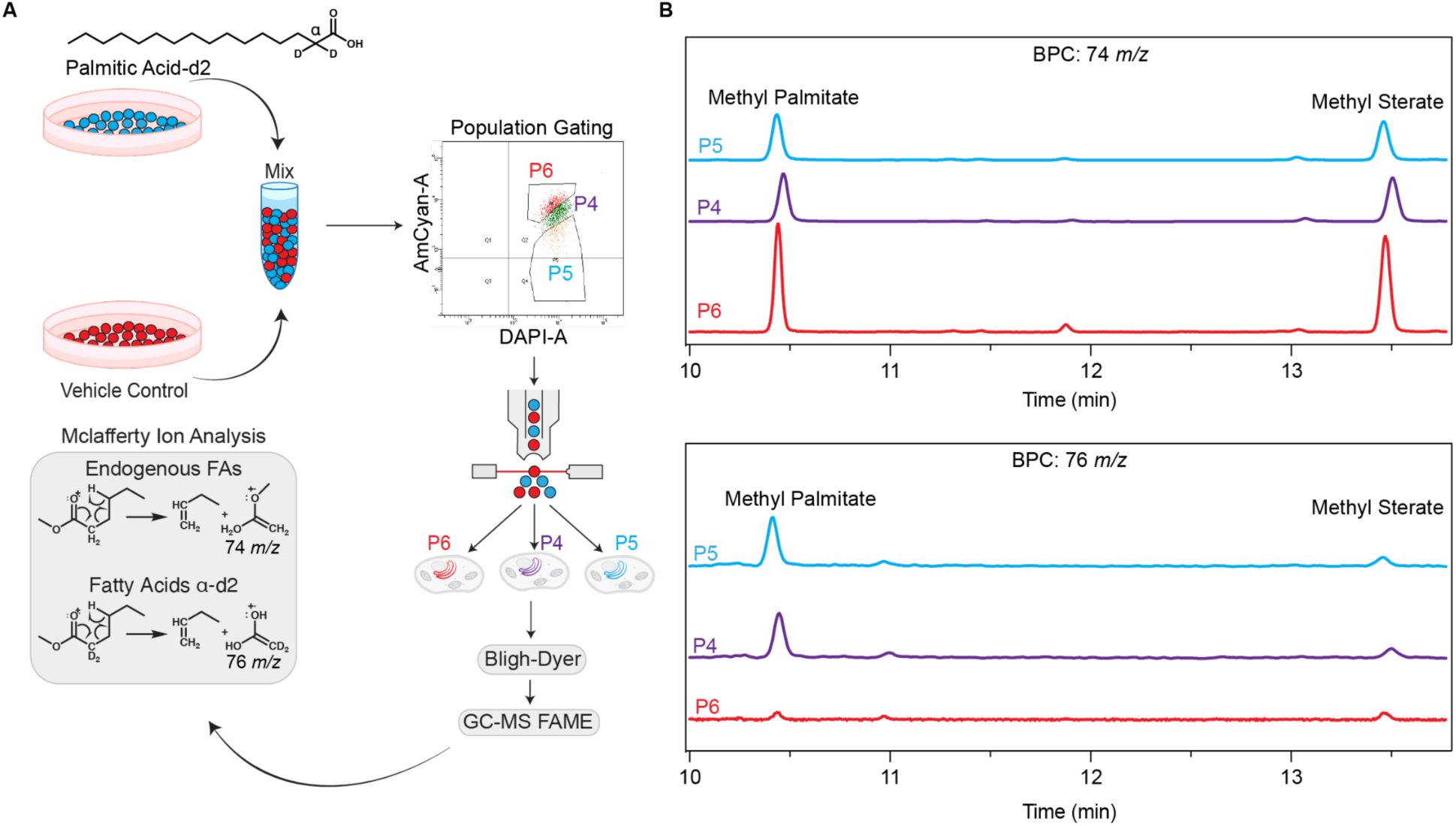
Sorting lipid-fed cells based on ER membrane fluidity. **A.** Cells treated with PA-d2 or vehicle control were mixed then sorted using a comparable DAPI-A filter for the ordered channel and AmCyan-A filter for the disordered channel. Cells were sorted into three groups: P6 (high fluidity, 216k cells), P5 (low fluidity, 86k cells) and P4 (intermediate fluidity, P4, 180k cells). Fatty acid profiles from the sorted cells were assessed by GC-MS FAME analysis. Confirmation that the P5 group were primarily PA-d2 treated cells was performed by assessment of the Mclafferty Ion shift from a *m/z* of 74 in endogenous fatty acids to 76 for the fatty acids with deuterations occurring on the α carbon. **B.** Extracted chromatograms found the 76 *m/z* ion species in the methyl palmitate peak in high abundance for the P5 (ordered) population and in moderate amounts in the P4 (intermediate) population. The 76 *m/z* ion species was also found in methyl stearate in the P4 (intermediate) population and the P5 (ordered) population. The 76 *m/z* ion was in low abundance in the high ER membrane fluidity P6 population, consistent with low saturated fatty acid incorporation in these cells.

## Summary

Here, we demonstrated that conjugation of small chemical moieties to C-Laurdan localizes the dye to internal organelles while retaining the spectral resolution needed to differentiate changes in lipid composition. We characterized multiple features of membrane packing allowed by the increased specificity provided by OTLs. We report robust apparent heterogeneity in the GP of each organelle of the four organelles targeted, likely reflecting regional differences in metabolic function. Measuring effects of chemical inhibitors supported proposed models of cholesterol trafficking as an important regulator of membrane fluidity in the lysosome, and the activity of the ETC affecting membrane packing in the IMM. Assessment of organelle-specific tolerances to palmitate treatment revealed OTLs could detect changes in fluidity due to change due to exogenous PA, a commonly used cellular model for metabolic disorders. We demonstrate that the sensitivity of ER-Laurdan is sufficient to sort cells that are PA-treated based on the resulting properties of their ER membranes. Together, these experiments demonstrate that OTLs are a versatile tool for probing intracellular membrane packing and its responses to lipidic modulators.

While chemical targeting of solvatochromic dyes is powerful for interrogating organelle-specific membrane responses, there are limitations that must be considered when interpreting results both between and within organelles. The OTLs reported here vary somewhat in their solvatochromic properties compared to Laurdan and C-Laurdan, with ER- and Mito-Laurdans showing diminished sensitivity to changes in lipid packing in LUVs compared to Lyso- and Golgi-Laurdans (Figure S4). Therefore, direct comparison of absolute organelle membrane fluidity between dyes must be considered carefully. For example, given the lower sensitivity of ER-Laurdan, the change in its GP with 200 µM PA treatment likely reflects a more dramatic relative sensitivity of ER membrane fluidity than what is reflected in the measurement. The solvatochromic sensitivity of these dyes could be improved through changes to the targeting moieties, e.g. *N*-tosylethylenediamines or propyl-chlorides (*38*, *59*) for ER or use of a longer linker to TPP. Alternatively, FLIM analysis could enhance sensitivity of all probes (*31*), though would also limit modalities, like cytometry, that currently rely on purely spectral measurements.

Despite the differences in spectral resolution, we demonstrate that OTLs are useful tools for measuring the regulators of membrane fluidity within each organelle. The cellular and physiological regulators of membrane fluidity regulators are vast and interconnected, given the numerous lipidic levers that regulate this property. Solvatochromic dyes like Laurdan can integrate these factors and provide a fluorescence measurement that is amenable to high resolution imaging of single cells or high throughput screening of populations. Organelle targeting these dyes provide clearer readouts for these applications, as changes in measurements using non-targeted probes can be driven by alterations to organelle abundances or proximities. By providing spatial sensitivity, the probes described here provide a new tool to untangle the metabolic regulation of cell membrane composition and physicochemical properties.

## Supporting information

Combined Supplemental Information

## Acknowledgements

The authors acknowledge Zulfiqar Mohamedshah, Shane Douty, and Dr. Neal Devaraj for helpful discussions and assistance with compound characterization, the UCSD Molecular Mass Spectrometry Facility for assistance with mass spectrometry, and the Human Embryonic Stem Cell Core for assistance with cell sorting. The work was supported by grants from the National Institutes of Health (NIH) (R35-GM142960) and the National Science Foundation (NSF) (MCB-2046303) to I.B. A.W. was supported by the NSF Graduate Research Fellowship Program (DGE-2038238) and the NIH Chemistry and Biology Interface Training Grant (5T32-GM146648).

## Methods and Materials

### Synthesis of *N*-(32-amino-3,6,9,12,15,18,21,24,27,30-decaoxadotriacontyl)-2,3,4,5,6-pentafluoro benzamide

BocHn-PEG_10_-NH_2_ (30mg) was dissolved in 1 mL of dry DCM with 89.6 mg Cs_2_CO_3_ and cooled to 0 °C. 63.25 mg 2,3,4,5,6-Pentafluorobenzoyl chloride (5.5 equivalents) was added dropwise and the mixture was stirred under N_2_ at 0 °C for 1 hour. The mixture was then stirred for an additional hour at room temperature before being dried *in vacuo*. 15 µL TFA in 1mL DCM was then added to neutralize the Cs_2_CO_3_ then filtered and dried *in vacuo*. Immediately after, 2 mL of a 1:1 DCM:TFA mixture was added and stirred for two hours under N_2_ at room temperature. The resulting deprotected PFP-PEG_10_ amine (Compound 1) was then purified by Preparative HPLC. All prep HPLC was performed on a 1260 Infinity II LC System equipped with a Agilent C18 column with a mobile phase of 0.1% formic acid in water and acetonitrile. NMR spectra were obtained on a 400 Varian Mercury Plus spectrometer at 21°C. ^1^H NMR (400 MHz, CDCl_3_) 3.75 (t, *J* = 4.9 Hz, 2H), 3.68 (t, *J* = 4.1 Hz, 6H), 3.62 (d, *J* = 4.5 Hz, 36H), 3.47 (s, 2H), 3.18 (q, *J* = 5.2 Hz, 3H).^13^C NMR (101 MHz, CDCl_3_) δ 160.50, 147.74, 145.22, 139.83, 112.16, 77.44, 77.32, 77.12, 76.80, 70.13, 70.10, 70.02, 69.96, 69.92, 69.88, 69.85, 69.82, 69.79, 69.68, 69.60, 69.45, 69.39, 66.90, 39.85, 39.72. Mass spectra were acquired on an Agilent 1260 Infinity Binary LC coupled to a 6230 Accurate-Mass TOFMS system. HR-ESI-TOFMS, *m/z* [M + H]^+^ calcd for 695.3173 found at 695.3167 *m/z*. Yield 67% as a clear oil.

### Synthesis of N-(5-(2-((6-dodecanoylnaphthalen-2-yl) (methyl) amino) acetamido)-3,6,9,12,15,18,21,24,27,30-decaoxadotriacontyl)-2,3,4,5,6-pentafluoro benzamide (ER-Laurdan)

2.0 mg of C-Laurdan (Tocris Bioscience) was dissolved in 300µL DMF along with 2.1 mg (1.1eq) of HATU, 3.84 mg DIPEA (6eq) and 0.34 HoBt mg (1.05 eq). The reaction was stirred under N_2_ for 5 minutes before 3.7 mg (1.05 eq) of compound 1 was added. The reaction was left stirring at RT temperature overnight. The reaction was then dried *in vacuo* and purified by Preparative HPLC (Compound 2). ^1^H NMR (400 MHz, CDCl_3_) δ 8.34 (s, 1H), 7.95 (dd, *J* = 8.5, 1.7 Hz, 1H), 7.83 (d, *J* = 9.0 Hz, 1H), 7.67 (d, *J* = 8.7 Hz, 1H), 7.10 (dd, *J* = 9.0, 2.5 Hz, 1H), 7.06 (s, 1H), 6.95 (s, 1H), 4.06 (s, 2H), 3.60 (h, *J* = 8.5 Hz, 36H), 3.49 (d, *J* = 4.0 Hz, 8H), 3.47 – 3.42 (m, 6H), 3.39 – 3.32 (m, 2H), 3.18 (s, 3H), 3.04 (t, *J* = 7.5 Hz, 2H), 1.82 – 1.70 (m, 2H), 1.47 – 1.34 (m, 2H), 1.26 (d, *J* = 6.7 Hz, 14H), 0.87 (t, *J* = 6.7 Hz, 3H). ^13^C NMR (101 MHz, CDCl_3_) δ 200.78, 170.59 149.24, 148.92, 145.35, 142.94, 137.60, 132.00, 131.48, 130.03, 127.01, 126.51, 125.36, 116.63, 114.49, 107.12, 77.80, 77.68, 77.48, 77.16, 70.68, 70.56, 70.51, 70.03, 58.33, 40.43, 40.41, 39.50, 38.94, 32.37, 30.11, 30.09, 30.01, 29.95, 29.81, 25.18, 23.15, 14.60. HR-ESI-TOFMS, *m/z* [M + H]^+^ calcd for 1074.5684 found at 1074.5678 *m/z*.. Yield 71% as a yellow oil.

### Synthesis of Golgi-Laurdan

2.0 mg of C-Laurdan was dissolved in 300 µL DMF along with 2.1 mg (1.01eq) of HATU, 5.18 mg DIPEA (8eq) and 0.34 HoBt mg (1.05 eq). The reaction was stirred under N_2_ for 5 minutes before 2.1 mg (1.05 eq) of the (*E*)-2-aminododec-4-ene-1,3-diol (d12:1 sphingosine) (Cayman) was added. The reaction was left stirring at RT temperature for 18 hours then dried *in vacuo* and purified by Preparative HPLC (Compound 3). ^1^H NMR (400 MHz, CDCl_3_) δ 8.35 (s, *J* = 1.8 Hz, 1H), 7.97 (d, *J* = 8.7, 1.7 Hz, 1H), 7.85 (d, *J* = 9.1 Hz, 1H), 7.69 (d, *J* = 8.7 Hz, 1H), 7.26 (s, 3H), 7.17 – 7.10 (m, 1H), 6.99 (s, 1H), 5.67 (dt, *J* = 14.2, 6.7 Hz, 1H), 5.42 (dd, *J* = 6.5 Hz, 1H), 4.26 (t, *J* = 5.4 Hz, 1H), 4.08 (s, 2H), 3.92 (ddd, *J* = 20.6, 9.2, 3.9 Hz, 2H), 3.70 (d, *J* = 3.3 Hz, 1H), 3.49 (s, 3H), 3.20 (s, 3H), 3.04 (t, *J* = 7.5 Hz, 2H), 1.91 (d, *J* = 7.1 Hz, 2H), 1.76 (q, *J* = 7.4 Hz, 2H), 1.52 – 1.18 (m, 24H), 0.87 (td, *J* = 6.9, 2.9 Hz, 6H). ^13^C NMR (101 MHz, CDCl_3_) δ 200.41, 170.92, 148.77, 137.17, 134.91, 131.88, 131.26, 129.69, 128.24, 126.73, 126.37, 125.14, 116.30, 107.13, 77.43, 77.31, 77.11, 76.79, 74.07, 62.28, 58.38, 54.83, 40.12, 38.59, 32.27, 32.01, 31.89, 29.75, 29.73, 29.64, 29.58, 29.45, 29.22, 29.19, 29.06, 24.82, 22.79, 22.74, 14.24, 14.20. HR-ESI-TOFMS, *m/z* [M + H]^+^ calcd for 595.4469 found at 595.4465 *m/z*. Yield 43% as a yellow oil.

### *Synthesis of (3-(2-((6-dodecanoylnaphthalen-2-yl) (methyl)amino)acetamido)propyl) triphenyl phosphonium* (Mito-Laurdan)

2.0 mg of C-Laurdan was dissolved in 300 µL DMF along with 2.1 mg (1.1eq) of HATU, 3.84 mg DIPEA (6eq) and 0.34 mg HoBt (1.05 eq). The reaction was stirred under N_2_ for 5 minutes before 2.1 mg (1.05 eq) of (3-aminopropyl)(triphenyl)phosphonium bromide (OOI Chemical) was added. The reaction was left stirring at RT temperature overnight. The following morning the reaction was dried *in vacuo* and purified by Preparative HPLC (Compound 4). ^1^H NMR (400 MHz, CDCl_3_) δ 8.09 (d, *J* = 1.8 Hz, 1H), 7.79 – 7.72 (m, 2H), 7.69 (dd, *J* = 8.7, 1.7 Hz, 1H), 7.56 (td, *J* = 7.8, 3.1 Hz, 6H), 7.43 (d, *J* = 8.7 Hz, 1H), 7.36 (dd, *J* = 12.8, 7.6 Hz, 5H), 7.26 (s, 3H), 7.18 (dd, *J* = 9.1, 2.6 Hz, 1H), 6.90 (s, 1H), 4.14 (s, 2H), 3.63 (d, *J* = 6.5 Hz, 1H), 3.50 (d, *J* = 6.1 Hz, 2H), 3.29 (s, 3H), 2.96 (q, *J* = 6.9 Hz, 2H), 2.33 (t, *J* = 7.6 Hz, 1H), 1.86 – 1.61 (m, 4H), 1.39 (td, *J* = 14.7, 6.5 Hz, 2H), 1.26 (d, *J* = 7.4 Hz, 14H), 0.87 (t, *J* = 6.7 Hz, 3H). ^13^C NMR (101 MHz, CDCl_3_) δ 200.27, 171.64, 148.92, 137.35, 135.30, 135.27, 133.23, 133.13, 130.80, 130.77, 130.61, 130.48, 129.71, 126.25, 125.41, 124.67, 118.14, 117.27, 116.35, 105.76, 77.43, 77.32, 77.11, 76.80, 57.08, 53.98, 40.66, 39.03, 38.86, 38.49, 32.01, 29.76, 29.73, 29.65, 29.61, 29.45, 24.90, 22.79, 22.23, 20.25, 19.72, 18.63, 17.50, 14.24. HR-ESI-TOFMS, *m/z* [M + H]^+^ calcd for 699.4074 found at 699.4064 *m/z*. Yield 62% as yellow oil.

### Synthesis of (3-(2-((6-dodecanoylnaphthalen-2-yl) (methyl)amino)-N-(3-morpholinopropyl) acetamide (Lyso-Laurdan)

2.0 mg of C-Laurdan was dissolved in 300 µL DMF along with 2.1 mg(1.1eq) of HATU, 3.84 mg DIPEA (6eq) and 0.34 HoBt mg (1.05 eq). The reaction was stirred under N_2_ for 5 minutes before 0.71 mg (1.05 eq) of 3-morpholinopropylamine (Sigma-Aldrich Inc) was added. The reaction was left stirring at RT temperature overnight, then dried *in vacuo* and purified by Preparative HPLC (Compound 5). ^1^H NMR (400 MHz, CDCl_3_) δ 8.32 (d, *J* = 1.6 Hz, 1H), 7.94 (dd, *J* = 8.7, 1.7 Hz, 1H), 7.83 (d, *J* = 9.1 Hz, 1H), 7.65 (d, *J* = 8.7 Hz, 1H), 7.38 (t, *J* = 5.9 Hz, 1H), 7.13 (dd, *J* = 9.2, 2.5 Hz, 1H), 6.88 (s, 1H), 4.08 (s, 2H), 3.73 (d, *J* = 8.2 Hz, 4H), 3.49 (s, 3H), 3.37 (d, *J* = 6.2 Hz, 2H), 3.03 (t, *J* = 7.5 Hz, 2H), 2.69 (t, *J* = 7.5 Hz, 2H), 2.44 (q, *J* = 9.3 Hz, 2H), 1.94 (t, *J* = 7.0 Hz, 2H), 1.39 (dd, *J* = 15.6, 7.1 Hz, 2H), 1.26 (d, *J* = 6.8 Hz, 14H), 0.87 (t, *J* = 6.7 Hz, 3H). ^13^C NMR (101 MHz, CDCl_3_) δ 200.27, 170.92, 148.92, 137.35, 135.30, 135.27, 133.23, 133.13, 130.80, 130.77, 130.61, 130.48, 129.71, 126.25, 125.41, 124.67, 118.14, 117.27, 116.35, 105.76, 77.43, 77.32, 77.11, 76.80, 63.50, 57.08, 53.98, 40.66, 39.03, 38.86, 38.49, 32.01, 29.76, 29.73, 29.65, 29.61, 29.45, 24.90, 22.79, 22.23, 20.25, 19.72, 18.63, 17.50, 14.24. HR-ESI-TOFMS, *m/z* [M + H]^+^ calcd for 524.3847 found at 524.3840 *m/z*. Yield 44% as a yellow oil.

### Preparation of vesicles and assessment of OTL spectral properties

LUVs comprised of either 100 mol % 1,2-dioleyol-sn-glycerol-3-phosphatidylcholine (DOPC) (Avanti), 55 mol % 1-palmitoyl-2-oleoyl-glycero-3-phosphatidylcholine (POPC) (Avanti) + 45 mole % cholesterol (Chol), or 55 mol % sphingomyelin (SM) (Avanti) + 45 mol % Chol, were prepared by evaporating a solution of lipids with N2 to a thin film then placing *in vacuo* for 1 hour to remove residual solvent. The solution was resuspended in HEPES (20 mM, pH 7.4) to a 1mM concentration and left rotating overnight. The following day, the solution was flash frozen and thawed 5 times, then extruded using a Mini-Extruder extruder (Avanti) through a 0.2 µm filter for 23 passages. Vesicles were stained with 5 µM of each Laurdan and left to incubate for 5 minutes. Spectral properties were acquired using a Cary Eclipse Fluorescence spectrometer equipped with a Cary Temperature Controller. Vesicles were allowed to equilibrate to their respective temperature for 5 minutes prior to acquisitions. For absorbance scans the excitation slit width was set to 1.5 nm and the emission slit width was 10 nm with absorbance being measured at 480 nm. For emission scans, OTLs were excited at 352 nm with a 10 nm slit width and absorbance 1.5 nm slit width.

### Cell culture and staining

HeLa cells were cultured in DMEM (Gibco), containing 25 mM D-glucose, 4 mM L-glutamine, and 1 mM sodium pyruvate, and supplemented with 10% FBS (Gibco) and 1% penicillin-streptomycin. Cells were incubated at 37 °C with 5% CO_2_ in a humidified atmosphere. Cells were routinely tested for mycoplasma by MycoStrip^TM^ (InvivoGen).

For labeling, cells were seeded on an 8-well chamber slides (LabTech) at 3x10^5^ cells per well for experiments performed 24 hours post seeding, and 2x10^5^ cells per well for experiments performed 48 hours post seeding. Cells were washed once with HBSS (Gibco) prior to being stained with ER-Laurdan (5 µM), Lyso-Laurdan (2.5 µM) and Mito-Laurdan (2.5 µM) for 20 minutes in Fluorobrite^TM^ (Gibco). All OTLs were dissolved in DMSO before being diluted to their respective concentrations in Fluorobrite. The staining solution was aspirated, and cells were then washed once with HBSS, then three times with Fluorobrite containing 10% FBS for six minutes per wash. Cells were then directly imaged in Fluorobrite. Cells stained with Golgi-Laurdan were washed once 24 hours prior to imaging then a 10 µM solution of Golgi-Laurdan in Fluorobrite containing 0.1% Fatty Acid Free BSA (Sigma-Aldrich) chilled to 4 °C was added to the cells and left to incubate at 4°C for 1 hour. Cells were then washed once with HBSS before the media was replaced. Prior to imaging cells were washed once with HBSS and imaged in Fluorobrite.

### Colocalization analysis

Colocalization experiments were performed with the following organelle markers post Laurdan staining for ER, Lysosomes and Mitochondria, respectively: ER-Tracker Red^TM^ (1 µM) (Thermo Fisher), LysoTracker^TM^ Deep Red (1 µM) (Thermo Fisher), MitoTracker Deep Red (500 nM) (Cell Signaling). To assess Golgi-Laurdan localization, cells were transfected with a SiT-mApple plasmid using Lipofectamine 3000 (Thermo Fisher). Briefly, 12 hours post seeding, media was aspirated and washed with HBSS and incubated in OptiMEM (Thermo Fisher) for 15 minutes. Lipofectamine 3000 plasmid mixtures were added, and cells were incubated for 4 hours. Media was replaced before being stained with the Golgi-Laurdan as described above.

### Cell treatments

For chemical inhibitor experiments, cells were seeded 24 hours prior to the experiment as described above. For treatment with U18666A (Sigma-Aldrich), cells were seeded in DMEM containing 10µM of the inhibitor then stained with Lyso-Laurdan the following day. For respiration uncoupler experiments, cells were stained with Mito-Laurdan as previously described, then Fluorobrite containing either 5 µM CCCP (Fisher Scientific) and 2.5 µM Rotenone (Sigma-Aldrich) was added and left for 15 minutes before imaging immediately after.

For PA and OA treatment, sodium palmitate (Thermo Fisher) or sodium oleate (Thermo Fisher) was complexed to fatty acid free BSA. The requisite fatty acid was dissolved in a 1:1 mixture of 0.1 mM NaOH in HBSS and ethanol to a 200 mM concentration and was heated to either 65°C for sodium palmitate or 40°C for sodium oleate. The fatty acid solution was added directly to a 4.4% BSA HBSS solution that had been heated to 37°C such that the final concentration of the fatty acid was 2 mM and at a 3:1 molar ratio with BSA. The mixture was stirred vigorously and incubated at 37°C for 1 hour. The fatty acid:BSA mixture or a BSA vehicle control was then diluted in DMEM or IMDM containing 10% FBS to the desired concentration and filtered through a 0.2 µM PES filter (Fisher Scientific) before use. Cells were incubated in their respective concentrations of the fatty acids for 24 or 48 hours prior to the experiment.

### Confocal microscopy

All microscopy experiments were done on a Zeiss LSM 880 equipped with a 63x/1.4 Plan-Apochromat oil objective. For colocalization assessment, OTLs were excited using a Diode 405nm laser line and emission was collected with an Airyscan detector using a 420-480 nm bandpass (BP); the corresponding organelle marker was excited with either a DPSS 561 nm or HeNe 633 nm laser lines and emission collected from a 560-620 nm BP or 645 nm longpass filters r. Images were acquired using ZEN Black and processed using default Airyscan settings.

For GP measurements, OTLs were excited with a Diode 405 nm laser and fluorescence was detected using a QUASAR GaAsP detector set to two spectrum windows: Ch1 (409-463 nm) for ordered channel detection and Ch2 (473-516 nm) for disordered channel detection. Due to the relatively disordered nature of internal organelles, Ch2 disordered detection was restricted to a shorter bandwidth than previously published (470-530 nm) bandwidths (*60*) to obtain signal intensity between the two channels in the same magnitude. Acquisition settings were kept constant between measurements for each OTLs. Individual cells were cropped for individual GP analysis. In each cropped image the GP was calculated as previously described (*32*, *60*). Briefly, the intensities of Ch1 and Ch2 were used to generate a binary mask to eliminate low signal pixels and the GP was calculated at each pixel by 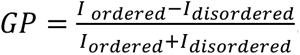.The G factor for GP normalization was determined by diluting each DMSO to the concentration used in cell medium, which was then imaged using the same acquisition settings. The G factor was calculated as previously described (*32*).

### Analysis of cells sorted using ER-Laurdan

HeLa cells were grown in a T25 flask to a density of 2x10^6^ in media supplemented with either 200 µM PA-d2 (Cayman) or vehicle control for two days. Prior to sorting the cells were stained as previously described, then treated with 5 mL accutase (STEMCELL) to gently detach cells. The cells were spun at 200xg for 10 minutes to pellet, then resuspended in Fluorobrite and mixed. Sorting experiments were done on an Aria II (BD Bioscience) cell sorter, equipped with a DAPI filter for ordered channel detection and a AmCyan filter for disordered channel detection. Cells were gated as shown in Figure S8 with the P5 and P6 gates differentiating cells emitting highly in the DAPI or AmCyan region compared to the main (intermediate) P4 population. Cells were pelted immediately after sorting and flash frozen until lipid extraction.

For FAME analysis, Cell pellets were resuspended in 100 µL H_2_O, then transferred to a glass tube containing 2 mL of 1:1 CHCl_3_:MeOH and vortexed. H_2_O (0.5 volume) was added and mixed by inversion, then spun at 1000 RPM for 5 minutes. The organic layer was collected and dried under N_2_ for 5 minutes. The film of lipids was then resuspended in 6 drops of toluene and 1.5 mL of 2.5% sodium methoxide in MeOH and heated to 50 °C for 30 minutes. The fatty acid methyl esters (FAMEs) were extracted in a 1:2 solution of hexane and MeOH. FAME samples were analyzed on an Agilent 8890 GC system equipped with a 60m DB25 column and a 5977B analyzer. The temperature was ramped from 40 to 230 °C over 20 minutes then held for 6 minutes as previously described (*11*). Samples were normalized to external FAME standards (Cayman). Data was acquired using Agilent MassHunter and analyzed using Agilent MassHunter Qualitative Analysis software.

### Statistical analyses

Significance analysis was determined using unpaired, two-tailed t-test using GraphPadPrism 10.2 software; p<0.05 was considered statistically significant. Population statistics for cytometry experiments were performed in FlowJo10.10.

