## Supplementary material for "Organelle-targeted Laurdans measure heterogeneity in subcellular membranes and their responses to saturated lipid stress": Combined Supplemental Information

This file contains:

- Supporting Figures S1-S8
- $^{1}H$ and $^{13}C$ NMR spectra for synthesized compounds

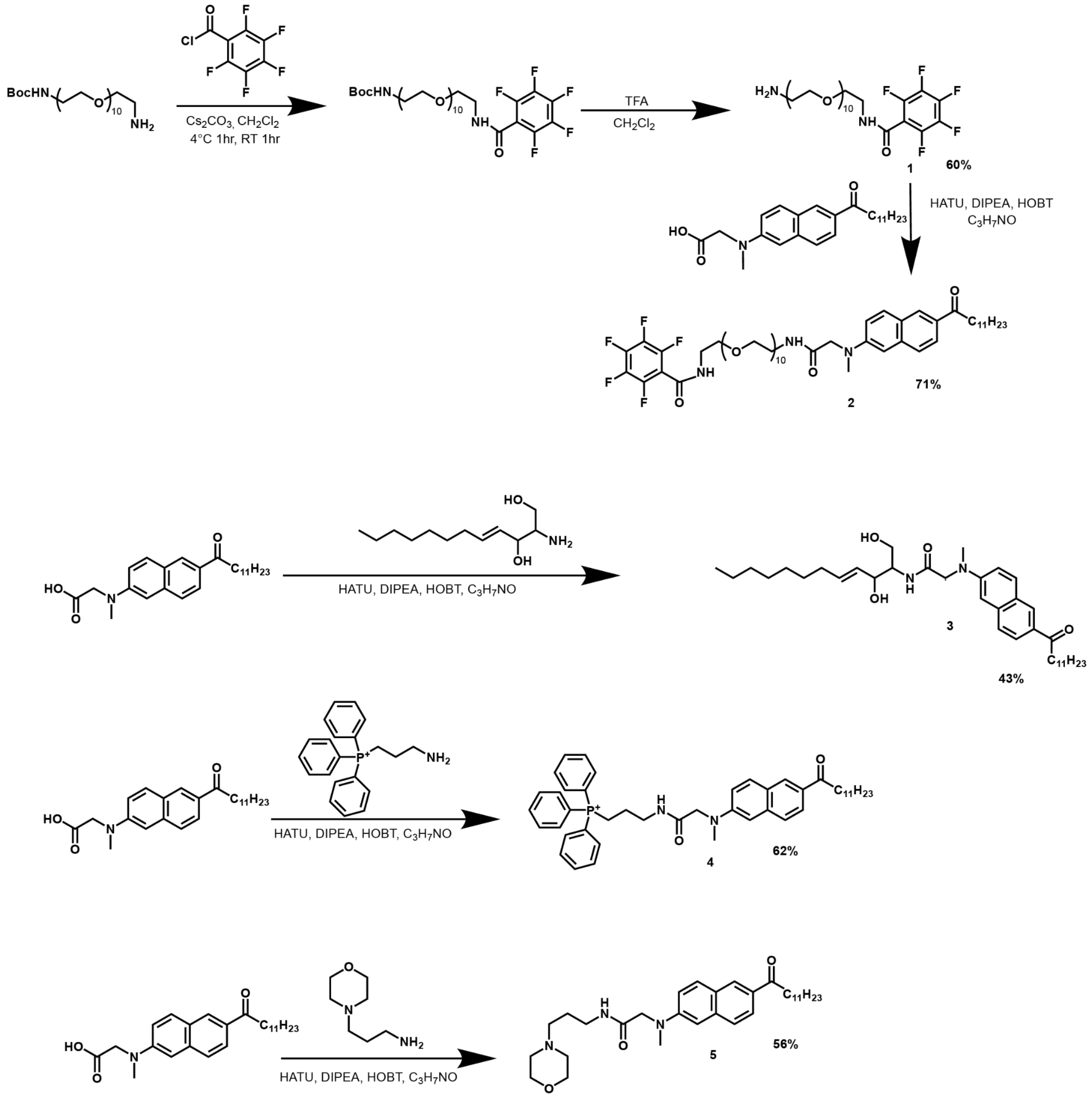


**Figure S1: Synthesis of Organelle-targeted Laurdans.** Synthesis schemes for OTLs. From top to bottom: ER-Laurdan, Golgi-Laurdan, Mito-Laurdan, Lyso-Laurdan. Details for each is available in the Materials and Methods.

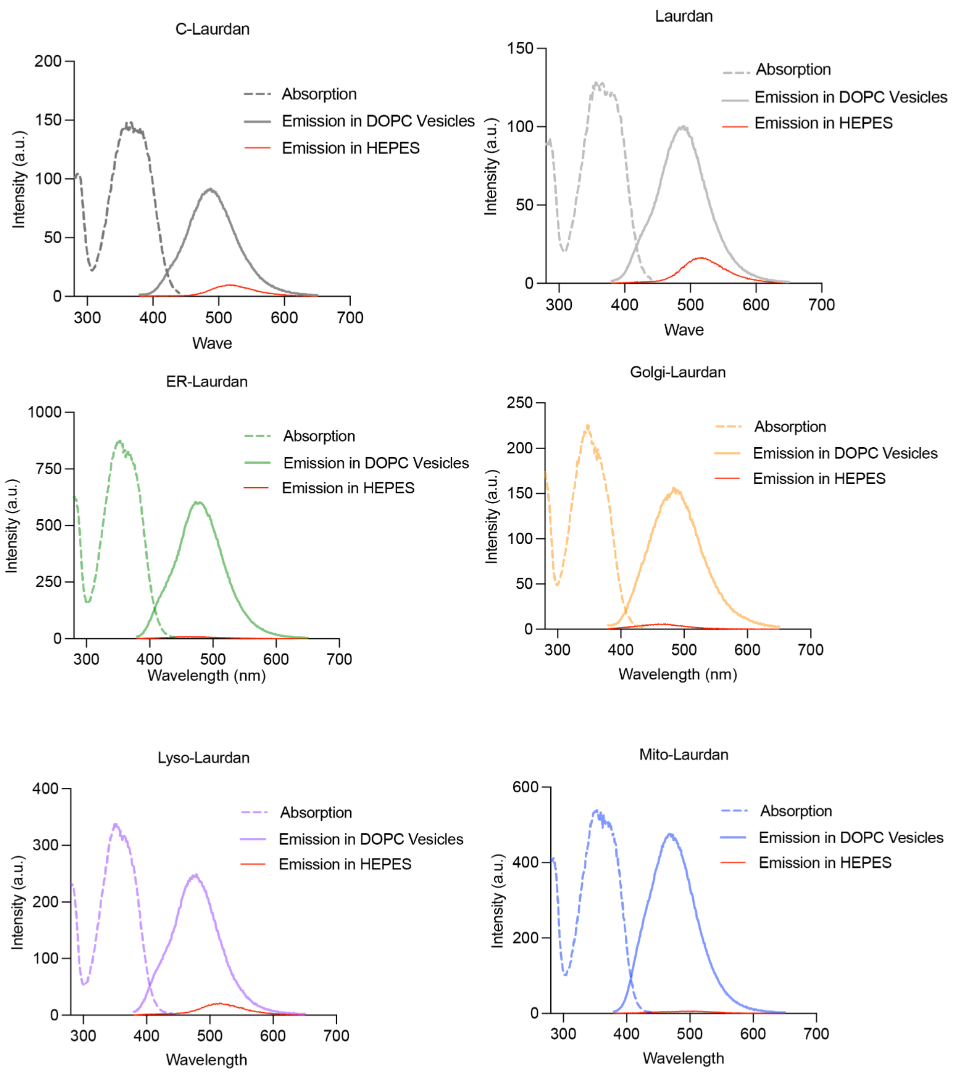


Absorption and Emission Maxima in DOPC Vesicles

| OTL | $\lambda_a^{max}$ | $\lambda_f^{max}$ |
|---|---|---|
| C-Laurdan | 360 | 488 |
| Laurdan | 356 | 490 |
| ER | 353 | 472 |
| Golgi | 347 | 483 |
| Lyso | 349 | 478 |
| Mito | 351 | 468 |

**Figure S2: OTLs retain absorption and solvent-dependency of the parent fluorophore**. Absorption and emission of Laurdans in DOPC vesicles. OTLs retain similar absorption (dashed lines) and emission (solid lines) maxima to Laurdan and C-Laurdan. Similar to the parent fluorophore, OTLs have minimal emission in aqueous solvents (red solid lines).

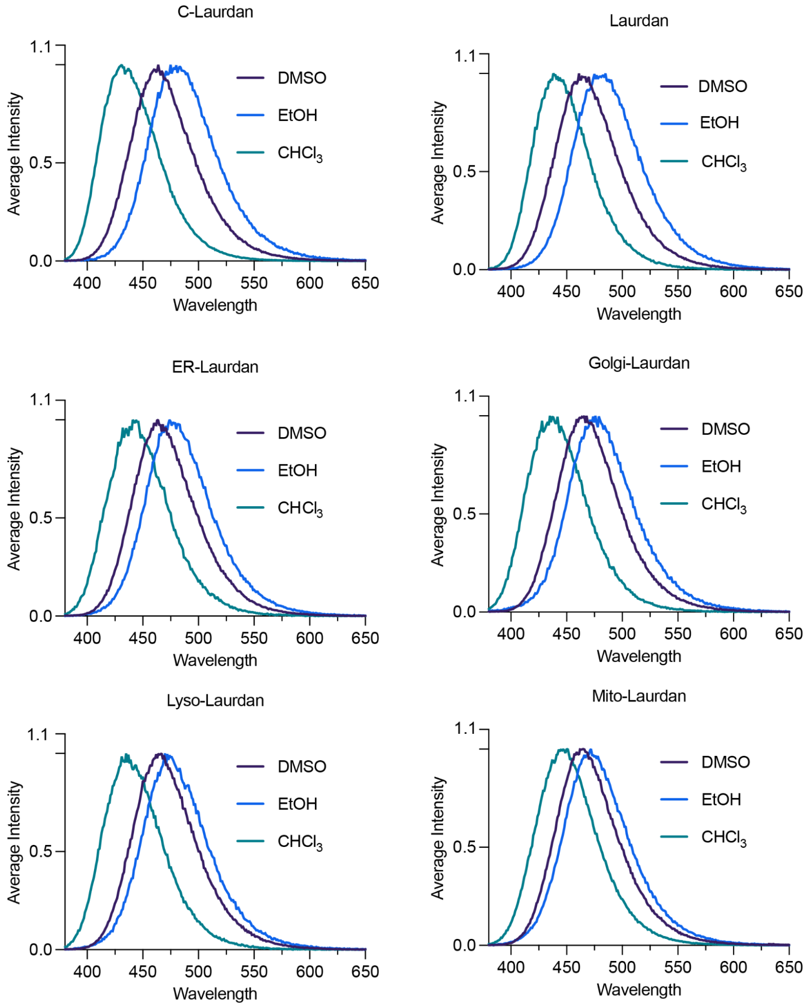


**Figure S3: Solvent-dependence emission of OTLs**. OTLs retain solvent dependent emission properties from the parent fluorophore. Emission spectra undergo a red shift with increasing solvent polarity ($CHCl_3$ < DMSO < EtOH).

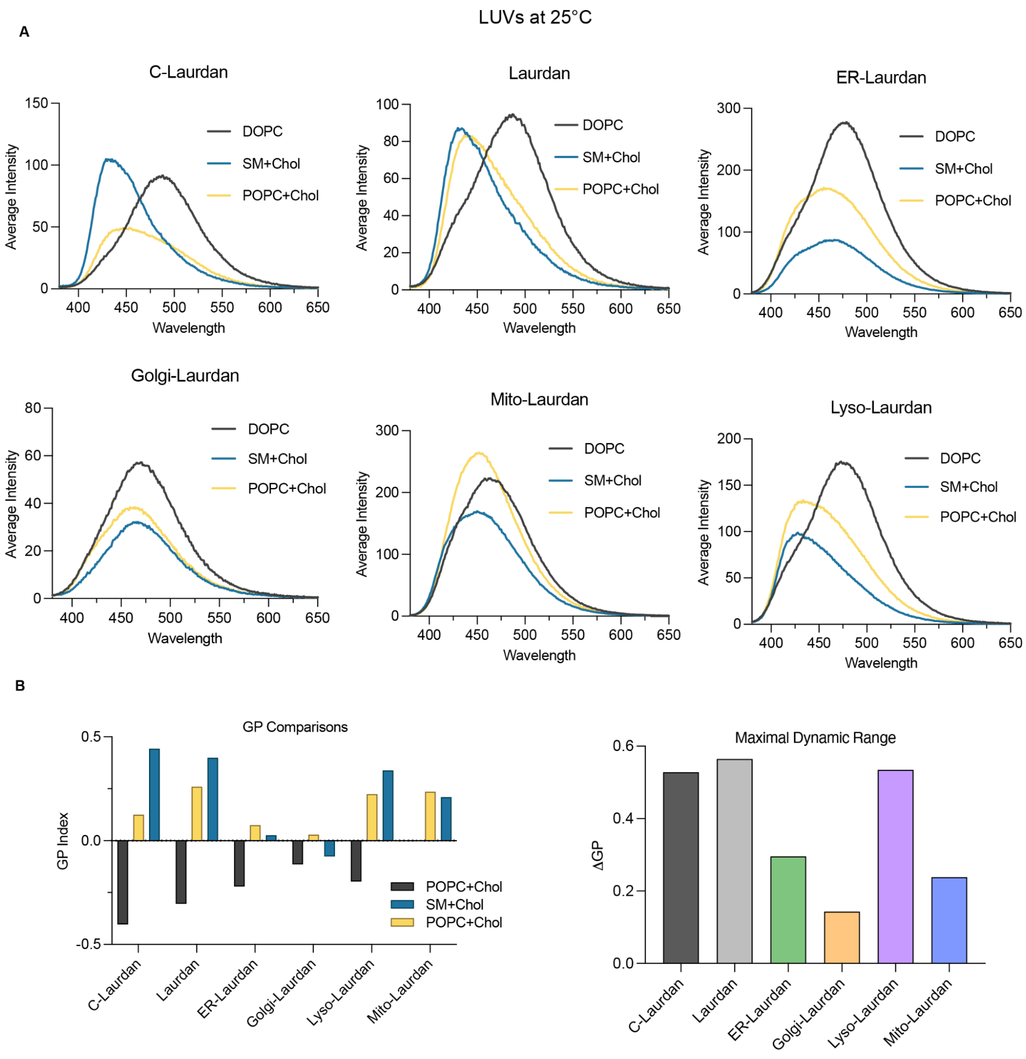


**Figure S4: Response of OTLs to lipid composition of synthetic vesicles at 25 °C**. **A**. Emission spectra of Laurdans in LUVs of varying fluidity at 25 °C. OTLs undergo a right shift to longer wavelengths in DOPC vesicles compared to SM+Chol and POPC+Chol. **B**. GP comparisons of Laurdans in LUVs. Emission intensity at 440 nm (ordered) and 490 nm (disordered) were used to calculate GP. All Laurdans show increases in GP in more ordered POPC+Chol and SM+Chol LUVs. **C**. Maximal dynamic range of Laurdans. OTLs show varying responsiveness to changes in vesicle composition. Maximal dynamic range was calculated by the difference between DOPC and SM+Chol GP (Lyso-, Golgi- and Mito-Laurdans) or POPC+Chol (ER-Laurdan). ER-, Golgi- and Mito-Laurdans show decreased sensitivity at 25 °C.

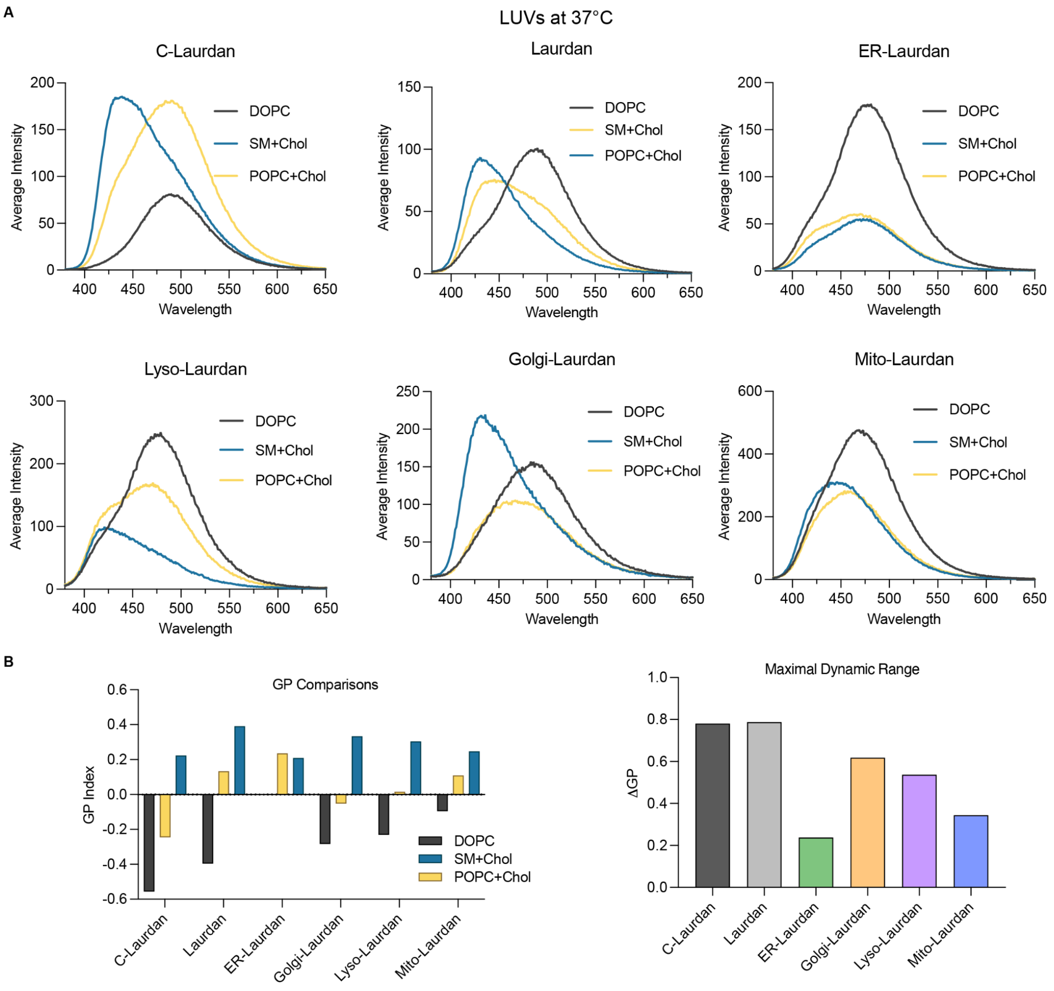


**Figure S5: Response of OTLs to lipid composition of synthetic vesicles at 37 °C**. **A**. Emission spectra of Laurdans at 37 °C. OTLs demonstrate a temperature dependence and show a more dramatic red shift in DOPC vesicles at physiological temperatures. A decrease in fluorescence intensity in ER-Laurdan stained SM+Chol and POPC+Chol compared to 25 °C suggests impaired integration of the dye into the LUV. **B**. GP comparisons between Laurdans show increased sensitivity to differences in membrane fluidity at 37 °C with the exception of ER-Laurdan which shows decreased responsiveness. **C**. All Laurdans show an increased dynamic range at 37 °C with the exception of ER-Laurdan.

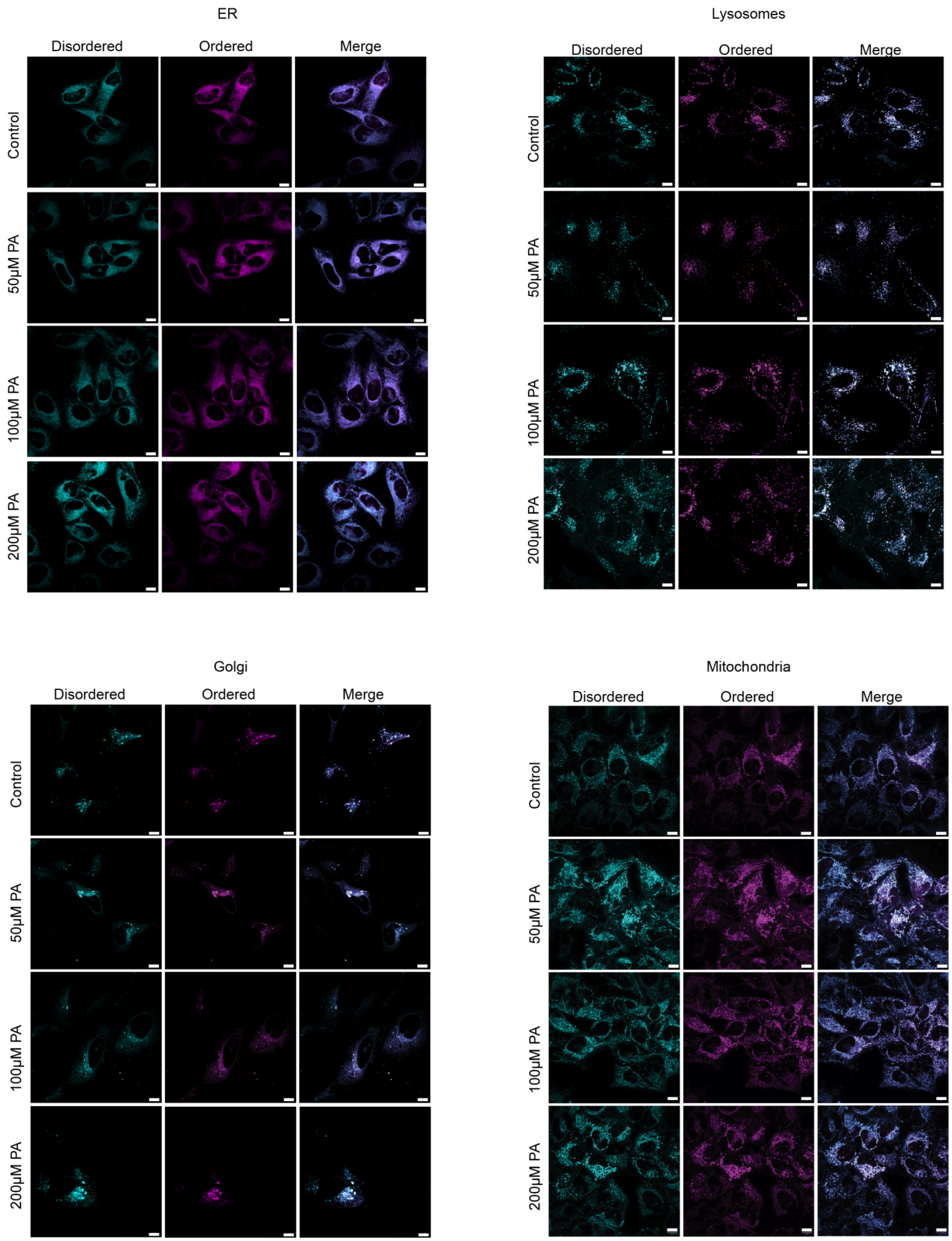


**Figure S6: Representative images from PA-treated cells not shown in Figure 5.** Representative images of palmitic acid treated cells used to calculate GPs in figure 5. Scale bars = 10 µm.

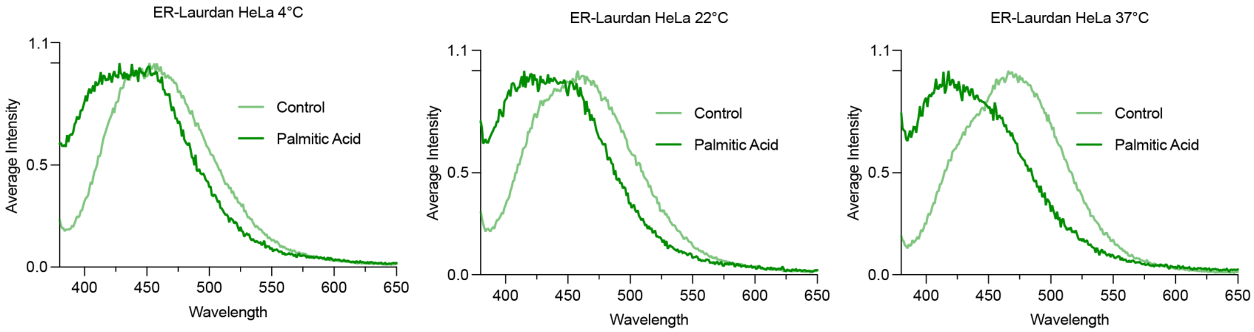


**Figure S7: Responses to PA treatment read out by ER-Laurdan in bulk cells**. HeLa cells (~$1.0 \times 10^6$) treated with 200 µM palmitic acid and stained with ER-Laurdan were trypsinized then the emission spectra was assessed using cuvette-based fluorometry. ER-Laurdan showed a more dramatic shift *in vivo* compared to what's observed in LUVs. In contrast to what's observed in liposomes, ER-Laurdan shows increased sensitivity at 37 °C to changes by PA treatment. The reduced sensitivity observed in liposomes could result from impaired integration of the unbound dye, since targeting is specific to free thiols in ER membranes.

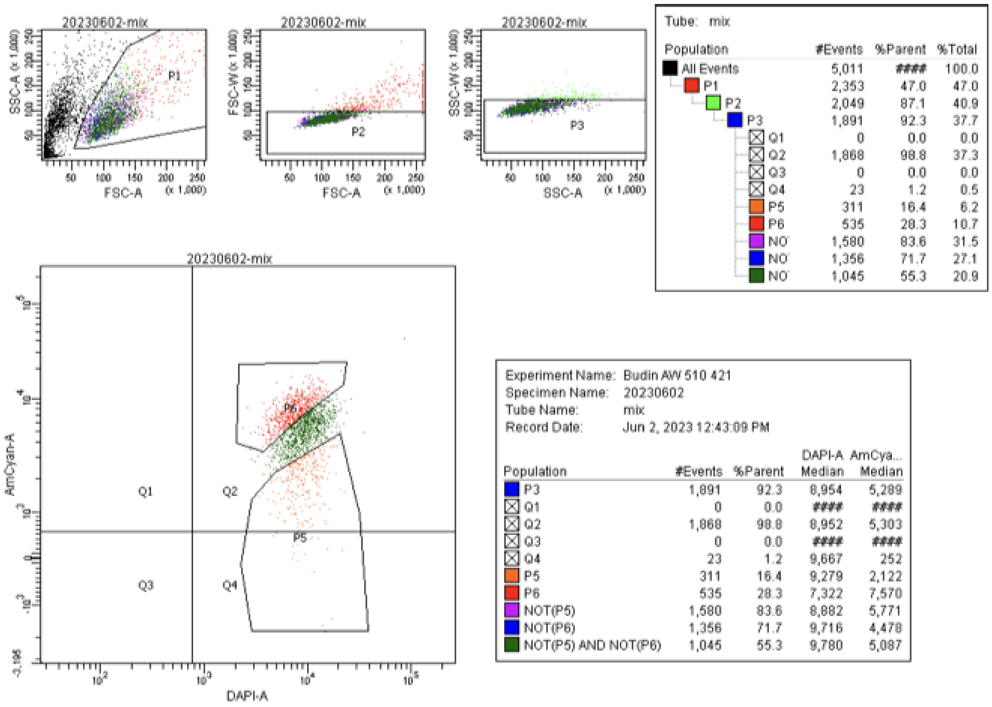


Tube: mix

| Population | #Events | %Parent | %Total |
|---|---|---|---|
| All Events | 5,011 | #### | 100.0 |
| P1 | 2,353 | 47.0 | 47.0 |
| P2 | 2,049 | 87.1 | 40.9 |
| P3 | 1,891 | 92.3 | 37.7 |
| Q1 | 0 | 0.0 | 0.0 |
| Q2 | 1,868 | 98.8 | 37.3 |
| Q3 | 0 | 0.0 | 0.0 |
| Q4 | 23 | 1.2 | 0.5 |
| P5 | 311 | 16.4 | 6.2 |
| P6 | 535 | 28.3 | 10.7 |
| NO | 1,580 | 83.6 | 31.5 |
| NO | 1,356 | 71.7 | 27.1 |
| NO | 1,045 | 55.3 | 20.9 |

Experiment Name: Budin AW 510 421
Specimen Name: 20230602
Tube Name: mix
Record Date: Jun 2, 2023 12:43:09 PM

| Population | #Events | %Parent | DAPI-A Median | AmCya... Median |
|---|---|---|---|---|
| P3 | 1,891 | 92.3 | 8,954 | 5,289 |
| Q1 | 0 | 0.0 | #### | #### |
| Q2 | 1,868 | 98.8 | 8,952 | 5,303 |
| Q3 | 0 | 0.0 | #### | #### |
| Q4 | 23 | 1.2 | 9,667 | 252 |
| P5 | 311 | 16.4 | 9,279 | 2,122 |
| P6 | 535 | 28.3 | 7,322 | 7,570 |
| NOT(P5) | 1,580 | 83.6 | 8,882 | 5,771 |
| NOT(P6) | 1,356 | 71.7 | 9,716 | 4,478 |
| NOT(P5) AND NOT(P6) | 1,045 | 55.3 | 9,780 | 5,087 |

**Figure S8: SI FACS Population gating**. Population gates from FACS experiment. Statistics of each population are shown on the table to the right. P5 and P6 gates were drawn in regions emitting highly in the DAPI and AmCyan regions, respectively. The P4 gate (intermediate) included all cells not collected in P5 or P6.
